# Fossils Do Not Substantially Improve, and May Even Harm, Estimates of Diversification Rate Heterogeneity

**DOI:** 10.1101/2021.11.06.467550

**Authors:** Jeremy M. Beaulieu, Brian C. O’Meara

**Affiliations:** Department of Biological Sciences, University of Arkansas, Fayetteville, Arkansas, 72701 USA; Department of Ecology and Evolutionary Biology, University of Tennessee, Knoxville, Tennessee, 37996-1610 USA

**Keywords:** fossilized birth-death, state speciation extinction, MiSSE, fossils, stratigraphic ranges, turnover rate

## Abstract

The fossilized birth-death (FBD) model is a naturally appealing way of directly incorporating fossil information when estimating diversification rates. However, an important yet often overlooked property of the original FBD derivation is that it distinguishes between two types of sampled lineages. Here we first discuss and demonstrate the impact of severely undersampling, and even not including fossils that represent samples of lineages that also had sampled descendants. We then explore the benefits of including fossils, generally, by implementing and then testing two-types of FBD models, including one that converts a fossil set into stratigraphic ranges, in more complex likelihood-based models that assume multiple rate classes across the tree. Under various simulation scenarios, including a scenario that exists far outside the set of models we evaluated, including fossils rarely outperforms analyses that exclude them altogether. At best, the inclusion of fossils improves precision but does not influence bias. Similarly, we found that converting the fossil set to stratigraphic ranges, which is one way to remedy the effects of undercounting the number of *k*-type fossils, results in turnover rates and extinction fraction estimates that are generally underestimated. While fossils remain essential for understanding diversification through time, in the specific case of understanding diversification given an existing, largely modern tree, they are not especially beneficial.

---

Diversification models, while fascinating to biologists, frequently lurch close to extinction themselves. For instance, Nee et al. (1994) demonstrated that estimating speciation and extinction rates from a molecular chronogram is theoretically possible. This was quickly followed by work from Kubo and Iwasa (1995), which showed that if rates varied through time there is an infinite number of alternative sets of time-varying speciation and/or extinction rates that produce the same number of lineages at any given point in time (an observation largely ignored at the time). State speciation and extinction models (-SSE; Maddison et al. 2007) were derived as a seemingly robust framework for estimating the direct effects of discrete traits on diversification rates. However, Rabosky and Goldberg (2015) found that if a tree evolved under a heterogeneous branching process, completely independent from the evolution of the focal character, SSE models will almost always return erroneously strong support for a model of state-dependent diversification. While this issue was partially rescued by the hidden state models of Beaulieu and O’Meara (2016), there remains some confusion as to whether SSE models remain a viable means of assessing state-dependent diversification (Rabosky and Goldberg, 2017; but see Caetano et al. 2018).

With Louca and Pennell (2020) comes a new salvo of criticism regarding diversification methods, which echo and expand on the earlier points by Kubo and Iwasa (1995), as well as past concerns about the possibility of estimating extinction rates generally (i.e., Rabosky, 2010). One postulated source of salvation has been the inclusion of fossils (Mitchell et al. 2019; Louca and Pennell, 2020; but see Černý et al. 2021). At a minimum, the inclusion of fossils should drastically reduce the number of congruent models by excluding any congruent model that assumes no extinction. We also know that fossils, even without formal models, have contributed greatly to the understanding of diversification processes — take the discovery of mass extinctions, for example. For these reasons, the fossilized birth-death (FBD) model (Stadler 2010) is seen as an appealing way of directly incorporating fossil information when estimating diversification rates, because it naturally assumes that the fossil information represents samples of extinct lineages in the past, in addition to the species sampled at the present. The purpose of this point of view is to investigate the improvement of estimation of diversification processes at just the tips of a tree.

## Importance of Sampling Ancestors

There are two main ways to incorporate fossils. The first is to use individual fossils placed as points on the tree (Fig. 1): they can be tips and/or along branches. The second is to represent stratigraphic ranges, which define the continuous duration of a species using only the oldest and youngest fossil appearance, ignoring how many fossils are sampled in between (Stadler et al. 2018). We first focus on the individual fossil process, as this has the most information, before examining the interval approach. An important, but frequently overlooked, property of FBD for individual fossils is that it distinguishes between two types of sampled lineages (Fig. 1). The first are referred to as *m* fossils, which represent sampled branches that went completely extinct before the present and did not give rise to any additional sampled lineages. The second type are referred to as *k* fossils, which represent samples of lineages that had sampled descendants (other fossils, or extant tips). In other words, these are fossils that represent ancestors sampled on internal branches that eventually lead to sampled species. The model assumes a sampling rate of fossils that does not vary across taxa or time — a questionable assumption, but probably no worse than similar simplifications that speciation rate or extinction rate is similarly invariant. However, while the model makes this distinction among these fossil types, the rate by which each is sampled is governed by the same global sampling rate, *ψ*, such that *ψ_m_* = *ψ_k_*.

**Figure 1.**
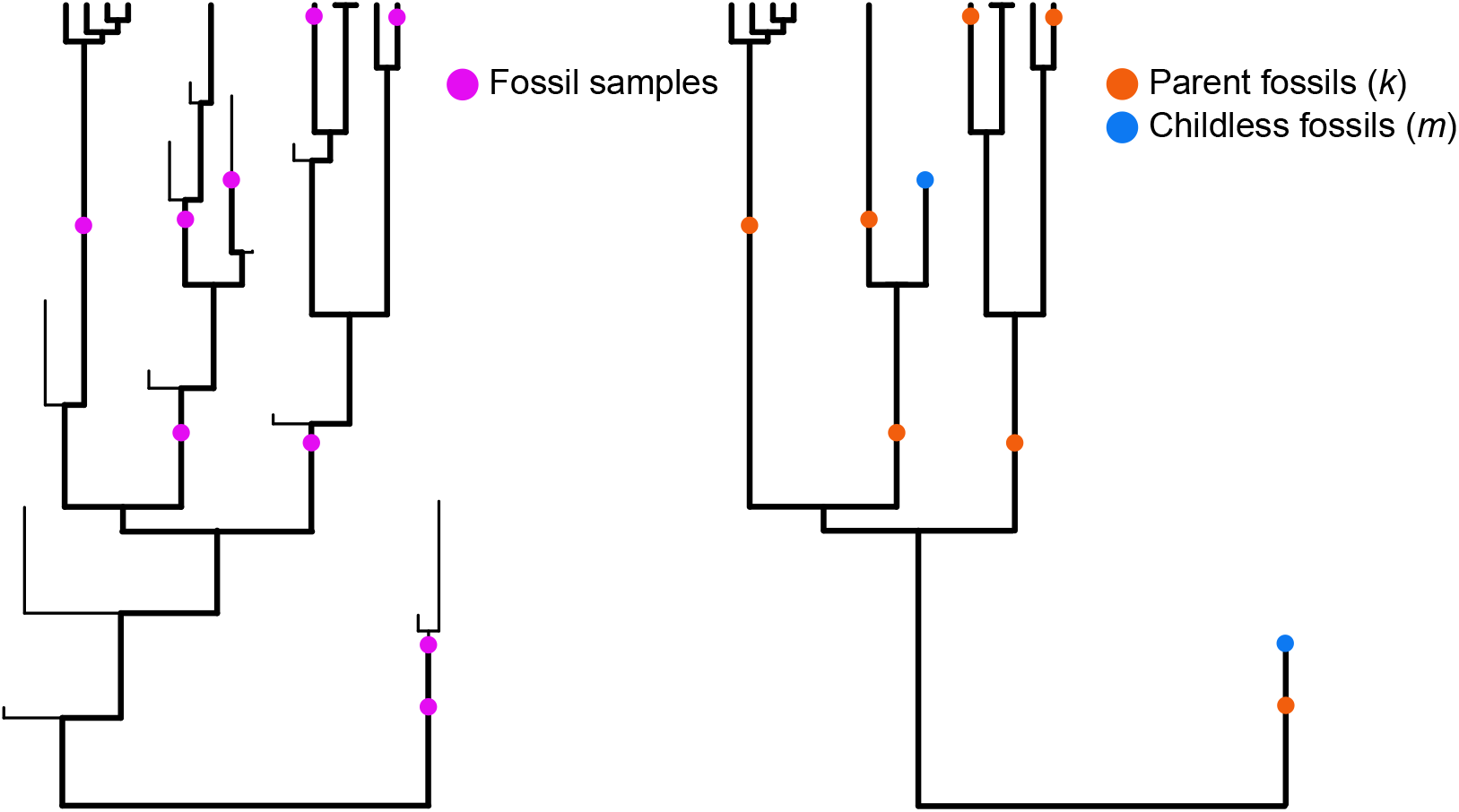
a) An example of a fossil set sampled from a complete tree generated by the birth death process, which includes both surviving and extinct members. b) The sampled phylogeny pruned down from (a) to only includes sampled tips and fossils. The fossilized birth-death model distinguishes between two types of sampled lineages. An *m* fossil (denoted by a blue dot) represents sampled branches that went completely extinct before the present and did not give rise to any additional sampled lineages (i.e., “childless” fossils). A *k* fossil (denoted by an orange dot) represents samples of lineages that had sampled descendants (other fossils, or extant tips; “parent” fossils).

The linking of these parameters makes sense. When a dinosaur gets washed into a river, the probability of it being excavated later by a paleontologist does not depend on whether a descendant seventy million years later is sampled (making our unlucky dinosaur a *k* fossil) or not (making this an *m* fossil). However, in the model, an assumption is not only that the probability of sampling the fossil is the same regardless of whether it has descendants or not, but also that the probability of including it in the phylogeny is the same whether or not it has descendants. For example, Paleobiology Database (PBDB; accessed October 2021) has 67 collections for *Tyrannosaurus rex* (Osborn 1905) and only eight for *Stegosaurus stenops* (Marsh 1887). For the FBD models to fully apply, there should be 67/8 times as many *T. rex* fossils as *S. stenops* fossils included in the tree. The FBD model of Stadler et al (2018), with its coarsening of the required data to only include stratigraphic ranges, reduces the need for not missing individual fossils but still makes assumptions of sampling of intervals.

Of course, there are other biases and variation in fossilization rate (the whole field of taphonomy studies this rich and varied process), but we are concerned with a potential bias of experimentalists undersampling *k* fossils. Even with approaches that infer whether a fossil is an *m*- or *k*-type from the data (Zhang et al. 2016), typically only one fossil per species is included (i.e., Ford and Benson 2020), and often infer zero *k* fossils on empirical data. This would have predictable effects on the estimates of *ψ* as well as the extinction rate itself, even if the methods place fossils perfectly. To better understand the dynamics of the different fossil types, here is Eq. (5) from Stadler (2010), which is probability density of a tree, *T*, with *n* extant taxa, *m* ≥ *0* fossils samples, and *n* ≥ *0* samples conditioned on the time since the most recent common ancestor of all taxa in the tree, *x_1_*.

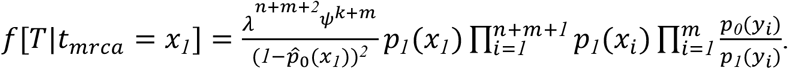

For specifics on what the probabilities *p_0_*(*t*), *p_1_*(*t*), and 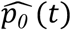 2(0 represent, we refer readers to the original derivation in Stadler (2010); for the purpose of this discussion, just note that they do not include *k*. We highlight two important observations here. The first is that the information provided by the *m*-type samples are used throughout the equation, including the time at which the extinct tip was sampled, *y_i_*, as well as the time at which they split from their common ancestor, *x_i_*. The second observation is that the only information provided by the *k*-type fossils is simply how numerous they are in the fossil set. In fact, it is irrelevant where exactly they occur on a branch or which branches have them — that is, the likelihood is the same if they are dispersed evenly across the tree or sampled in one 10,000-year interval on one edge. This point is best illustrated by rewriting the numerator in the first term in the equation above to separate out *k* and *m*,

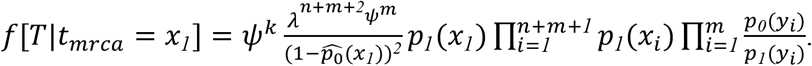

Moving the *ψ^k^* to the front of the equation shows that the effect that *k*-type samples have on the probability is simply based on a factor of *ψ^k^*. What this also says is that when both *m*- and *k*-type fossils are included, the overall log-likelihood can be calculated as, *logL* = *logL_m only_* + [*k* * *log*(*ψ*)]. As an example, take the tree and set of sampled extinct lineages presented in Figure 2d. The overall log-likelihood for the maximum likelihood estimate (MLE) for this tree is −1224.937. The log-likelihood assuming only *m*-type fossils, but using the same parameter estimates, is −983.6938, and we can add 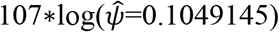 to obtain the overall log-likelihood of −1224.937 (for more details on these calculations see Supplemental Materials).

**Figure 2.**
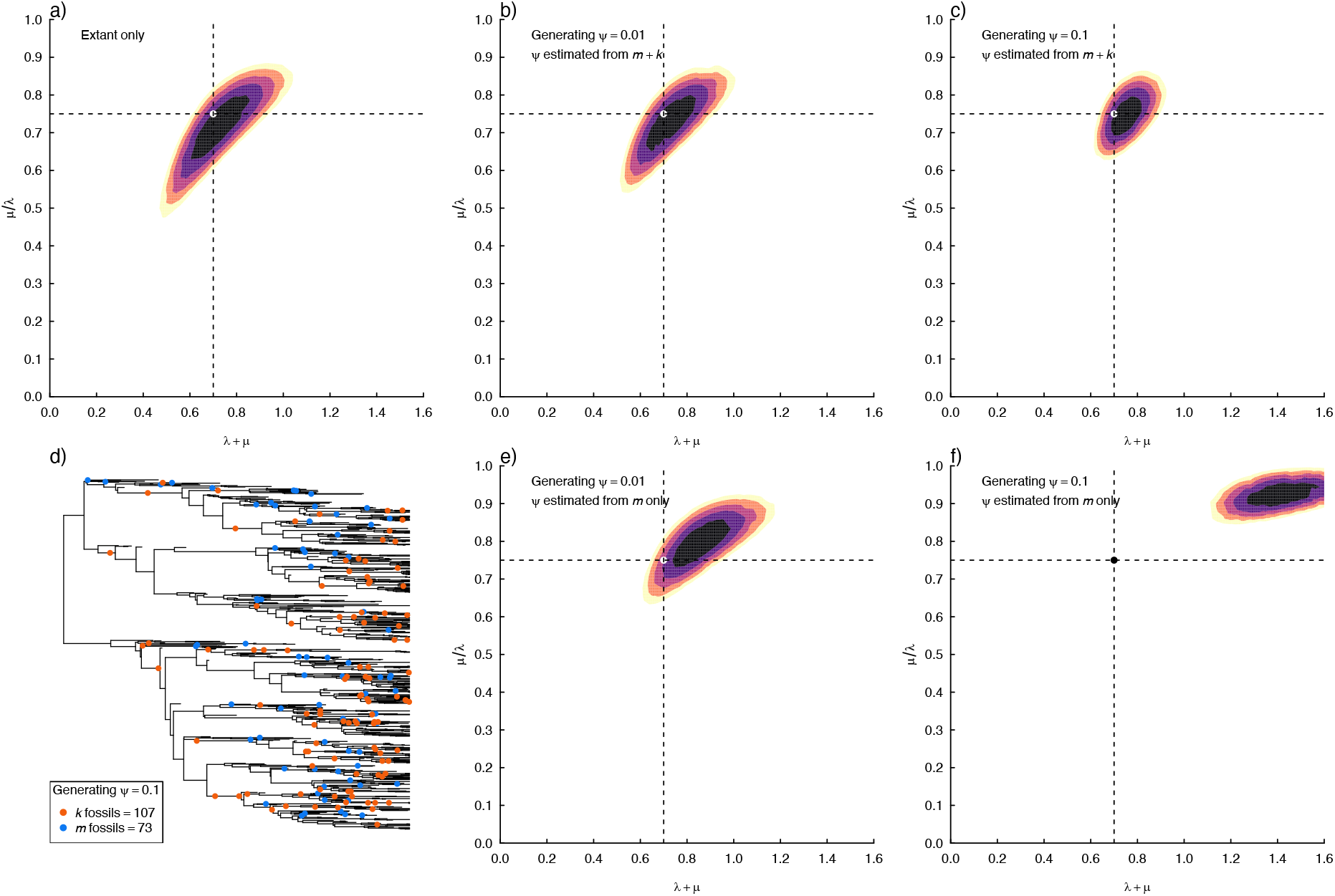
Contour plots of the likelihood surface under various scenarios for the same tree (topology shown in d) simulated under a birth-death process. Each surface is constrained such that turnover rate (*τ* = *λ* + *μ*) and extinction fraction (*ε* = *μ/λ*) are set given a pair of fixed values, but *ψ*, the rate of fossil preservation, is free to find their MLE. We sampled 5000 pairs of points for *τ* and *ε* from a latin hypercube sampling design. When provided with a sample of extinct lineages that is perfectly consistent with the generating model, the FDB performs and behaves well and generally reduces the variance in turnover rate and extinction fraction (b, c), relative to ignoring fossils completely (a). However, if *k* fossils are removed completely (e,f), the likelihood surface begins to erroneously shift away from the generating parameters towards regions of parameter space of very high extinction rates as values of *ψ* increase. The dashed vertical line represents the generating value for turnover rate (*τ*=0.70) and the dashed horizontal line represents the generating value for extinction fraction (*ε*=0.75).

On the one hand, if a tree was generated by a Yule process (i.e., no extinction), the inclusion of only *k*-type samples has no effect on the parameter estimates. Their inclusion will simply slide the overall log-likelihood down, again, by a factor *k*\*log(*ψ*). On the other hand, in the case of non-zero extinction rates, if *m*-type fossils are included and *k*-type not added, this will have a profound impact on the parameter estimates, particularly with regards to *ψ* and *μ*. Under the FBD formulation, the linking of *ψ_m_* = *ψ_k_* forces the model to interpret the lack of *k*-type fossils as evidence of a low sampling rate. This will be in tension with the presence of only *m*-type samples, such that to explain a low sampling rate, but with many *m*-type fossils in the set, the extinction rate must have been substantial. As shown in Figure 2e-f, this is exactly what happens when the *k*-type fossils are removed from the fossil sample set. The likelihood surface erroneously shifts away from the generating parameters towards regions of parameter space of very high extinction rates (based on estimates of turnover rate, *λ* + *μ*, and extinction fraction, *μ/λ*).

In any case, when provided with a sample of extinct lineages that is perfectly consistent with the generating model, the FDB performs and behaves exactly as it should (Fig. 2b-c), but so does a model that ignores fossils completely (Fig. 2a). Our concern, however, is what happens when theory meets practice. An important element of the FBD model is the possibility of having sampled ancestors, long a question in paleontology (see Foote 1996). There is a rich literature investigating the theoretical plausibility (Foote 1996; Gavryushkina et al. 2014) and statistical identifiability of ancestors in the fossil record (Gingerich 1979; Fisher 2008; Parins-Fukuchi et al 2019; Wagner 2019; Parins-Fukuchi 2021). Many of these predict that sampled ancestors may be common. For example, Zhang et al (2016) ran simulations that resulted in approximately half to three-quarters of the fossils being ancestral to descendant samples (other fossils and/or modern taxa). We simulated over a wide range of conditions (Fig. 3) and found that under most conditions, sampled ancestors should outnumber other kinds of fossils. To address a reviewer’s question about empirical findings, we examined a few papers that had used the FBD model for empirical data. Sampled ancestors outnumbering other kinds of fossils is not the pattern typically found with FBD studies. For example, Gavryushkina et al. (2017) found 18 of 35 fossils were ancestral, but Slater et al. (2017) and Pyron (2017) appeared to find fewer than 2% of fossils as being ancestral, based on nonzero terminal branch lengths of their main trees. However, we note that with FBD, if people are presenting consensus branch lengths as an average, then it is possible that most of the posterior probability is for zero length branches (thus an ancestral fossil), but with a handful of samples with the fossil not being ancestral to lead to a nonzero average branch length.

**Figure 3.**
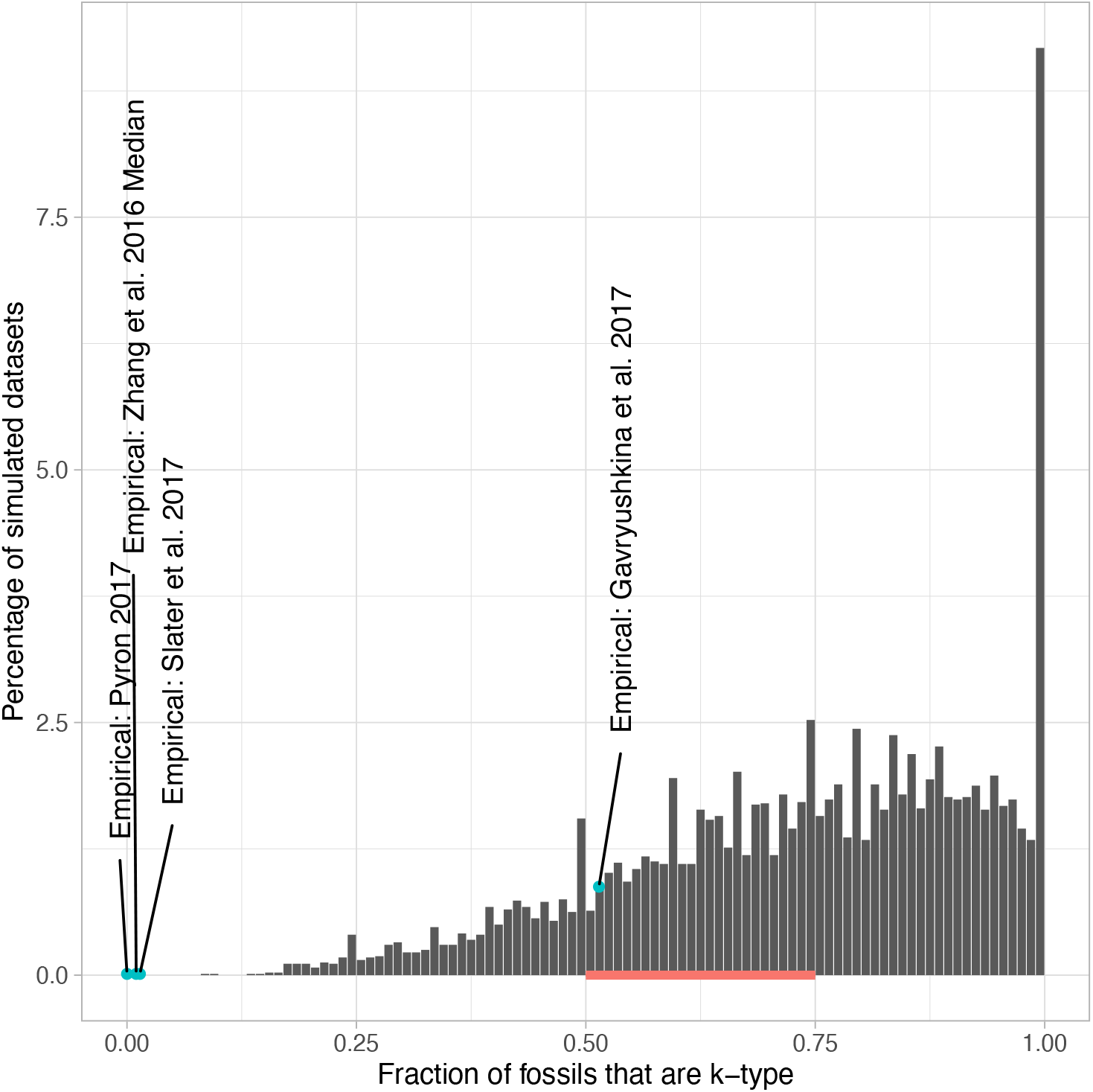
Histogram of the fraction of *k*-type fossils sampled across a large set of simulations over a wide range of conditions in relation to *k*-type fossils used in empirical FBD studies. Specifically, we generated a grid that make up all possible combinations of elements from vectors of turnover rate (incrementing *τ* = 0.1 to 1.0, sampling every 0.1), extinction fraction (incrementing *ε* = 0.05 to 0.95, sampling every 0.1), and sampling rate (*ψ* = 0.1, 0.05, 0.10, 0.20). We repeated each generating parameter 20 times, resulting in 8000 simulation replicates in total, with the histogram representing the distribution of the fraction of *k*-type fossils sampled across all replicates. Despite the wide range of simulation conditions, empirical studies (blue dots) appear to have fewer sampled ancestors. The red line along the x-axis represents the approximate range of simulated values from Zhang et al. (2016), and the blue dot corresponding to the study of Zhang et al. (2016) represents the median fraction across the range used in their analysis of Hymenoptera.

There are many other potential reasons for mismatch between empirical and simulated data. We focus here on just one possible explanation, undersampling of *k*-type fossils. The likelihood equations of Stadler et al. (2010) all assume that a dataset is not censored to only include *m*-type fossils, though some implementations of the FBD allow for using priors prohibiting sampled ancestors (see comparison conducted by Matzke and Wright, 2016). But, as far as we can ascertain, there is no conditioning for this censoring in the likelihood equations, and it may lull empiricists into believing that sampling ancestors is not a necessary requirement. Of course, as noted above, fossils need not only be treated as individual fossils placed as points on the tree. A recent extension of the FBD explicitly treats the same fossil sets above as multiple samples of the same species that specify distinct stratigraphic ranges (Stadler et al. 2018). All that is required, then, is the oldest and youngest fossil within a range, as opposed to the total number of fossils, which is then marginalized out, potentially rendering issues related to counting *k* fossils somewhat moot in most cases.

These are the topics we seek to investigate here. Specifically, we explore the benefits of including fossils in more complex models that assume multiple rate classes across the tree in comparison to excluding them altogether. In short, will fossils help diversification analyses largely based on modern phylogenies? Our hypotheses at the outset were that 1) well-sampled fossils would have a very beneficial effect on estimation of diversification parameters versus having no fossils and that models that exclude *k*-type fossils would perform poorly, likely being outcompeted by models with no fossils; 2) we expected approaches with stratigraphic samples would have less benefit (reduced data, especially about fossilization rate) but also would be relatively insensitive to sampling issues; and 3) we expected that fossils would be especially valuable in a situation where the true model was more complex than any model used in analysis, which approximates the situation in reality. To keep analyses feasible, we assumed that we knew the topology and branch lengths of the tree, and timing and placement of any fossils used, perfectly. Improvement of empirical trees from inclusion of fossil taxa, or harms coming from incorrect fossil placement or uncertain timings, are beyond the scope of this study.

## Fossilized *SSE

Incorporating the sampling rate, *ψ*, into a birth-death model that allows multiple discrete shifts has been implemented before (e.g., fossil BAMM, Mitchell et al. 2019). However, here we incorporate fossils within our hidden state speciation and extinction framework (HiSSE; Beaulieu and O’Meara, 2016), that includes any number of observed and/or hidden states. We focus our tests here exclusively using MiSSE (see Vasconcelos et al. 2021), a likelihood-based, hidden state only model, like BAMM, but without priors (which could help, hurt, or have little effect on inference), but also, importantly, without the discontinuous inheritance of extinction probability that made the likelihoods used in BAMM mathematically incorrect (see Moore et al. 2016). It is relatively straightforward to extend the equations of the canonical FBD model of Stadler (2010) to the state speciation and extinction models of Maddison et al. (2007) so that they also include both state transitions and the tree-wide rate of fossil sample rate parameter, *ψ*:

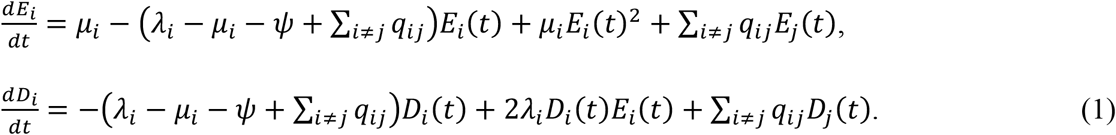

The probability *E_i_*(*t*) is the probability that a lineage starting at time *t* in state *i* leaves no descendants at the present day (*t*=0), and *D_i_*(*t*) is the probability of a lineage in state *i* at time *t* before the present (*t*>0) evolved the exact branching structure as observed. These ordinary differential equations are generalized so that any number of observed or hidden states can be included in the model. For character-based models, like HiSSE and MuHiSSE, *i* and *j* represent the different observed and hidden states combinations specified in the model, whereas with MiSSE *i* and *j* represent hidden states only. For an extant tip, the initial condition for *D_i_*(*0*) is *ρ_i_*, which defines the probability that an extant individual observed in state *i* is sampled in the tree and *1* – *ρ_i_* for *E_i_*(*0*). For an *m*-type fossil, *D_i_*(*t*)=*ψE_i_*(*t_e_*), and *E_i_*(*t*)=*E_i_*(*t_e_*), where *t_e_* represents the time at which the sampled extinct lineage was sampled; for *k*-type fossil, *D_i_*(*t*)=*ψD*(*t*), and *E_i_*(*t*)=*E_i_*(*t*). At nodes, *N*, the initial condition is the combined probability of its two descendant branches, *L* and *R*, such that, *D_N,i_* = *D_L,i_D_R,i_λ_i_*. The overall likelihood is the sum of *D_N,i_* calculated at the root.

When the fossil set represents stratigraphic ranges, the likelihood calculation is much more involved and requires designating three specific types of edge segments. First, we note that our implementation reflects the “symmetric speciation only” portion of Stadler et al. (2018; see Corollary 13), where only bifurcating speciation events are allowed — that is, a speciation event reflects the extinction of an ancestral species, and two new descendant species arise. This greatly reduces the complexity of the original model formulation, which at its most complex, also includes both anagenetic (ancestral species is replaced by a new descendant) and asymmetric (a single descendant “buds” off a persistent ancestral lineage) speciation processes with their own rates. Second, we note that a stratigraphic range is defined only by the oldest (*o*) and youngest (*y*) fossils that are unequivocally assigned to a species across a distinct time interval. Thus, a stratigraphic interval records the known duration of species on branches. These ranges are represented by a single fossil and a tip (i.e., fossil is *y*=0), by a branch fossil and an extinct tip (*y*=*t_e_*), by two fossils representing a sampled ancestral stratigraphic range along an edge (*o*>*y*), or even by a single fossil (i.e., *o*=*y*). Fossils that occur within a given range are ignored.

When an edge segment is not associated with a stratigraphic range, the branch calculation and starting conditions proceed exactly as described in Eq. 1 above. However, when an edge segment represents a stratigraphic range, we must modify *D_i_*(*t*) to remove the possibility of an unobserved speciation and subsequent extinction event within the interval given that we know that the entire segment [*o,y*] belongs to the same species:

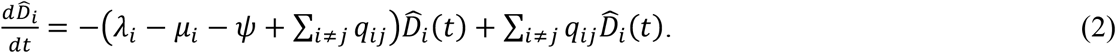

When *y*=0, such that it is an extant species, the initial condition for 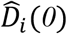 is *ρ_i_*; if *y* is an *m*-type fossil, 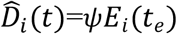; if *y* denotes the beginning of a sampled ancestor stratigraphic range then 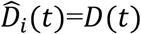. Finally, for edge segments that represent intervening time intervals between two stratigraphic ranges without an observed speciation event, we modify the branch calculation to account for zero or more unobserved speciation events. Again, given the assumption that a stratigraphic range represents the sampling of a single species across a distinct interval of time, the presence of two stratigraphic ranges along the same edge would imply that at least one unobserved speciation event had occurred somewhere within this interval. To account for this, at the rootward end, *y_a_*, which represents the youngest fossil of the older of the older of the two stratigraphic ranges, we correct *D_i_*(*t*) following the Stadler et al. (2018):

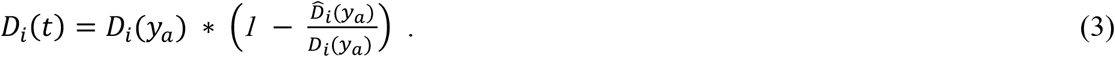

This probability then becomes the starting condition for the next stratigraphic range. Note that none of three specific branch types requires altering *E_i_*(*t*) from what is presented in Eq. 1, because this probability is based solely on time and is therefore unaffected by the tree topology. As before, at nodes, *N*, the initial condition is the combined probability of its two descendant branches, *L* and *R*, such that, *D_N,i_* = *D_L,i_D_R,i_λ_i_*. The overall likelihood is, again, the sum of *D_N,i_* obtained at the root, but we must also marginalize over the number of fossils within a stratigraphic range. This is done by multiplying the sum of *D_N,i_* at the root by *ψ^K′^* and *e^ψL_s_^*, where *K’* represents the total number of sampled fossils that represent the start and end times of a stratigraphic range (if *o*=*y* for a given range then this counts as only one fossil), and where *L_s_* represents the sum of all stratigraphic range lengths 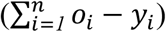.

For both the original FBD, and FBD with stratigraphic ranges, we condition the overall likelihood by *λ_i_* [*1* – *E_i_*(*t* | *ψ* = *0*)]*^2^*, where and *E_i_*(*t* | *ψ* = *0*) is the probability that a descendant lineage of the root survived to the present and was sampled assuming no sampling in the past. We note that fossil BAMM does not condition on survival in this way, or even at all, when the tree includes extant taxa. For character-based models, such as HiSSE and MuHiSSE, we weight the overall likelihood by the probability that each possible state gave rise to the observed data (see FitzJohn et al. 2009). However, it was pointed out by Herrera-Alsina et al. (2018) that at the root, the individual likelihoods for each possible state should be conditioned prior to averaging the individual likelihoods across states. It is unclear to us which procedure is correct, but it does seem that both weighting schemes behave exactly as they should in the case of character-independent diversification — that is, the overall likelihood reduces to the likelihood of the tree multiplied by the likelihood of a trait model. We have also tested the behavior of both (not shown), and the likelihood differences are very small, and the parameter estimates in simulation are nearly indistinguishable from one another.

In the absence of character information, assuming a model with a single birth rate and death rate, and a *ψ*>0 the likelihoods are identical to that of Stadler (2010) and Stadler et al. (2018). To ensure that our math and the implementations are correct with two or more birth and death rates, we relied on the property of SSE models described by Beaulieu and O’Meara (2016) and Caetano et al. (2018): when a trait has no differential effect on the diversification process (i.e., the character-independent model), the overall likelihood is the product of the tree likelihood and likelihood of a Markov model applied to the character data (it is the sum of the two likelihoods in log space). Thus, for a binary character that is independent of the diversification process on a tree with two rate classes and a non-zero fossil preservation rate, the BiSSE likelihood should reduce to the product of the likelihood of a MiSSE model with two rate classes and the likelihood of the transition rate model applied in the character data. As we show in the Supplemental Materials, we can confirm that our implementation of the FBD for MiSSE, HiSSE, and MuHiSSE (two binary characters; Nakov et al. 2019) are indeed correct. We also show that the different branch calculations for FBD with stratigraphic ranges conforms to the analytical calculations provided by Stadler et al. (2018). All models are available in the R package *hisse* (Beaulieu and O’Meara 2016).

## How Do Fossils Help in a World Where All Models are Simply Wrong?

Mitchell et al (2019) showed that with their fossil implementation within BAMM, which allows for shifts in discrete regimes of speciation and extinction, there were substantial improvements to the rate estimates, most notably with the extinction fraction. These results seem at odds with our results presented thus far, as well as results from a set of additional single rate regime simulations where trees were generated using a range of speciation and extinction rates (see Table S1; Supplemental Materials). While their simulations were focused on understanding the behavior of a much more complex model, it should be noted that the most striking improvement in extinction fraction occurred when *ψ*=1 fossils/Myr, which is an extremely high rate of preservation. In our case, the maximum value *ψ*=0.1 fossils/Myr resulted in a fossil set that was at least as numerous as the number of extant taxa (*N*>200), which seemed reasonable to us. Still, even when the addition of fossils resulted in doubling the amount of data available to the birth-death model, there was very little impact on the rate estimates, especially when the generating extinction fraction was high (*ε* = 0.75; Figure 2). We also wondered how other functions of speciation and extinction, such as turnover rates and net diversification rates, are impacted not only by the inclusion of fossils, but also by the inclusion of biased samples. The simulations of Mitchell et al. (2019) demonstrated that estimates of speciation are insensitive to the inclusion of fossils, with extinction being strongly impacted, which suggests that this would also impact additional functions of speciation and extinction in different ways.

Similar to the single regime simulations above, when we first simulated under our MiSSE model under scenarios of moderate difficulty, the main effect of including fossils when estimating of functions of speciation and extinction (namely, the turnover rate, *τ_i_* = *λ_i_* + *μ_i_* and the extinction fraction, *ε_i_* = *μ_i_/λ_i_*) was, again, to simply reduce the variance of the estimates (i.e., calculated as the mean of the squared errors from the *x*_model-averaged_) across the different rate regimes, not the bias (i.e., calculated as the mean of *x*_model-averaged_ - *x*_true_; see Supplemental Materials for more details). As expected, removing *k* fossils completely from the set results in very high biases related to turnover rates specifically, as *ψ* increases, and rather severe downward biases generally in estimates of net diversification rates (*r_i_* = *λ_i_* – *μ_i_*). With regards to converting the fossil set to stratigraphic ranges, there was a curious and general tendency for a downward bias in both turnover rate and extinction fraction.

An easy and valid criticism of these types of simulations is that they are too simplistic. Trees were generated under an SSE model that shifted between, at most, two different rate classes, with the rate of these shifts set by a single transition rate, *q*. While we were genuinely surprised that extant-only trees performed as well as trees that also included fossil sample data under the same conditions, the simulation scenarios are hardly realistic. The processes that generate most empirical trees are likely very complex, likely carrying signatures of non-random extinction, even mass extinctions, with diversification rates varying substantially among lineages and across time. The simplicity of our MiSSE models is out of mathematical convenience and tractability. Even still, we might expect that even when the true model is not included in the set of models evaluated, the inclusion of fossil information attached to branches distributed across the tree would provide more weight and better parameter estimation for the more complex models that are included in the set. Most models will work fine with data that meet their assumptions, but with complex and messy data, maybe fossils are needed to rescue their performance.

To explore this further we devised a simulation scenario that was meant to closely imitate processes and conditions that are more realistic in empirical settings, which would also prove challenging to our MiSSE model, but also be biased in a way that would favor including fossils. Specifically, we simulated 100 trees where we assumed that there were four discrete “regimes” of turnover rates (*τ_A_* = 0.3, *τ_B_* = 0.6, *τ_C_* = 1.5, *τ_D_* = 1.0 events/Myr) and extinction fractions (*ε_A_*= 0.7, *ε_B_* = 0.9, *ε_C_* = 0.95, *ε_D_* = 0.8) that controls the diversification dynamics. We also assumed a rather extreme heterogeneous transition matrix, **Q**, that governed the dynamics of transitions, *q_ij_*, among these four regimes,

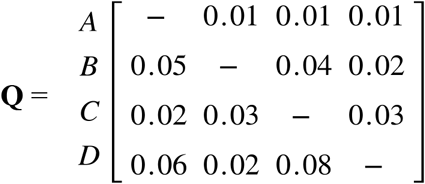

The magnitude of these rates was chosen somewhat at random, but cumulatively ensured that approximately 10% of the branches across the entire history of the tree (including extinct lineages) contained at least one change in rate regime. We also encoded two mass extinction events. The first occurred after the first 40-time units and removed, at random, 70% of lineages alive at that time point. The second occurred 30-time units later, 70-time units in total from the start of the simulation and removed 90% of the lineages alive at the time point (see Fig. 4a). Finally, simulations were terminated once the tree reached 267 taxa. However, we non-randomly chose 67 taxa and removed them from the final tree. The purpose of this procedure was to assume that our final tree had a biased sampling fraction of 75%. The bias was generated by simulating a trait under Brownian motion, then normalizing the values so that they were between 0 and 1 and had a phylogenetic signal. These values were then used as probabilities for removing taxa.

**Figure 4.**
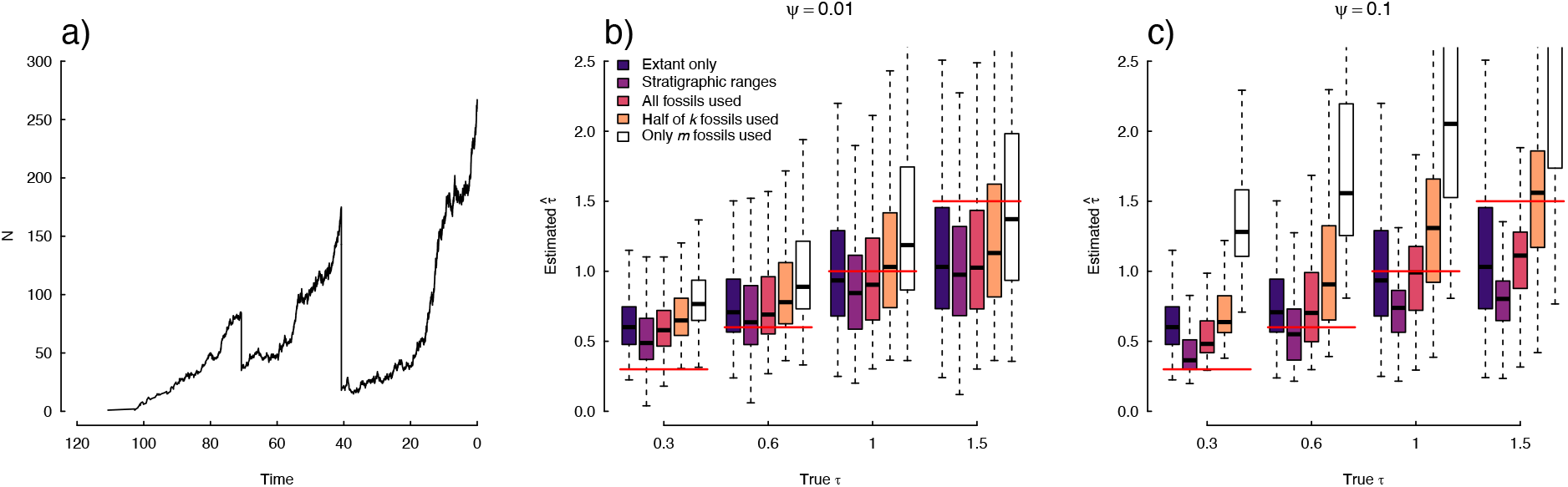
The uncertainty surrounding estimates of turnover rate when the models being evaluated does not include the true model. Specifically, we assumed that there were four discrete “regimes” of turnover rates (*τ_A_* = 0.3, *τ_B_* = 0.6, *τ_C_* = 1.5, *τ_D_* = 1.0) and extinction fractions (*ε_A_* = 0.7, *ε_B_* = 0.9, *ε_C_* = 0.95, *ε_D_* = 0.8) that controls the diversification dynamics; we also assumed a rather extreme transition dynamics among these four regimes (see text). We also encoded two mass extinction events as shown by the lineage through time plot (a), and we also assumed biased sampling among the tips that survived to the present. The boxplots represent the distribution of model-averaged rates based on the true rate classes at the tips. (b-c) Despite the true model being far outside the models being evaluated, the turnover rates at the tips hints at there being more complexity within the data than even the most complex two-rate models in the set might suggest. However, the inclusion of fossils does not seem to perform remarkably better than not including them.

We know fossilization rates vary, sometimes dramatically, by time and taxa (Wagner and Marcott 2013). Incorporating this would make the simulation far more realistic as well, but it would provide additional complications given the difficulties in separating variable extinction rates from variable preservation rates (e.g., Foote et al. 2019; Louca et al. 2021). The fact that the fossilization process matches that of the model gives the fossils a chance to perform better than not including them. That is, can well-modeled fossils help an analysis where the true diversification model is more complex and different than any analyzed? If fossils cannot help in this case, they probably will not help. Had we used a more complex fossilization process, and fossils failed to help in that case, it could just be that we tried too extreme a difference and set up a bias against the utility of fossils. In all cases, we assumed that all the models started with the true trees and that any fossils that were sampled were placed perfectly, both in time and on the tree. These assumptions are quite optimistic; our goal was to find the impact of fossils on diversification rate estimates, not assess the impact of fossils on all aspects of tree inference (which would require simulation of morphology and other traits, state-based fossilization rates, tree inference, homology assessment, and more), nor make it difficult for fossils by adding realistic issues of difficulty with placement, taphonomic biases, and more.

For each simulation replicate, we fit four MiSSE models where we set *f_i_* = 0.75 to account for incomplete sampling (though assuming, incorrectly, that all taxa have the same sampling rate; see Beaulieu 2020). Specifically, we fit a single regime model, a model that only allows turnover rate to vary (*τ_A_* ≠ *τ_B_*), a model that only allowed extinction fraction to vary, and a model that allowed both to vary (*τ_A_* ≠ *τ_B_* and *ε_A_* ≠ *ε_B_*). Each model was evaluated using a two-step optimization routine. The first step consists of a bounded stochastic simulated annealing run for 5000 iterations, followed by a bounded subplex routine that searches parameter space until the maximum likelihood is found. This model set ensured that, as with real data, the set of models were far simpler than the true generating model. We also fit the same set of models for increasing values of *ψ* (i.e., 0.01, 0.05, 0.10 events/Myr), and for different sampling of *k*-type fossils (i.e., full, half *k*, only *m*-type fossils used), including converting the set to stratigraphic intervals. When converting a fossil set into a set of stratigraphic ranges, we simply take the oldest and youngest fossil on each edge, removing all others, prior to pruning unsampled extinct lineages from the full tree. We chose to summarize results based on the diversification rates model-averaged across only the tips that survived to the present, because the rate regimes in the generating model do not map to the rate regimes in the model set, namely, regime *D* in a four-rate model does not easily map to regime *A* or *B* in a two-rate model. For a given model, the marginal probability of each rate regime is obtained for every tip, and the rates for each regime are averaged together using the marginal probability as a weight: a weighted average of these rates is then obtained across all models using Akaike weights. The use of model-averaging is particularly advantageous here because the tip rates are not conditional on a single best model that would be far less complex than the model that generated the data. Different models reflect different aspects of the data. By fitting multiple models, each good model might tell us something about how the data evolved. Model-averaging is one way to summarize these different elements in a comprehensible way.

Despite the true generating model being far outside the set of models evaluated, the use of model-averaging at least hints at there being more complexity within the data than even the most complex two-rate models allow. When the model-averaged tip rates are aligned with the four true tip rate regimes for each simulation iteration there are consistent significant positive slopes across all treatments (Fig. 4). For fossil-based analyses, particularly for those where the number of fossils was greater than the number of extant tips (i.e., *ψ* = 0.10 fossils/Myr), there are clearer differences between the four rate regimes compared to the extant-only estimates. However, as with all other analyses presented here, when the fossil set is perfectly consistent with the FBD process by which the samples were taken, the main effect of the fossils is to reduce the overall uncertainty in the tip rate estimates compared to extant-only inferences.

There were some improvements with respect to the parameter estimates with the inclusion of the full fossil sets overall, most notably at two ends of the rate distribution under the generating model (i.e., *τ_A_* = 0.3 and *τ_C_* = 1.5 events/Myr). Incidentally, both rate classes were by far the most frequent states across the entire history of almost every simulated tree. The highest turnover rate regime also had the highest extinction fraction, and this coupled with several mass extinctions, resulted in very few survivors at the tips. Even with the addition of a substantial number of fossils placed throughout the tree, the improvement in inferring this particular rate category from model-average rates was minimal. From a qualitative standpoint, by ignoring biases in rate and focusing on comparisons among tips, it seems that all treatments, regardless of the fossil makeup, can consistently assign the correct sign differences among pairs of tip rates. That is, even when the trees were analyzed with only half of the *k*-type fossils used, if they were removed entirely, or converted to stratigraphic ranges, the expected proportion of sign differences are consistent with the extant-only and full fossil set estimates (Table S2). Not surprisingly, as the number of *k*-type fossils are removed, the model-averaged estimates of turnover rate become greatly inflated, from erroneously high estimates of the extinction, and are associated with much greater uncertainty. When the fossil sets are converted to stratigraphic ranges, the turnover rates are, again, generally underestimated, with the improvement of the estimates with the lower turnover rate regimes being simply incidental. Taken together, this is consistent with the MiSSE scenarios above (Fig. S1), as well as the constant birth-death models (Fig. 2e-f), which suggests that: 1) there is only a modest improvement in the rate estimates when perfectly placed fossils are included; 2) gross omissions of *k-*type samples in a FBD erroneously shifts estimates towards areas of very high turnover rates and extinction fraction; and 3) converting to stratigraphic intervals results in erroneous shifts towards lower turnover rates and extinction fraction.

## A Caution About Stratigraphic Ranges

Our simulation results highlight a curious pattern in that when a fossil set is converted to stratigraphic ranges, estimates of turnover rate and extinction fraction are increasingly underestimated as the generating *ψ* increases. A clear illustration of the pattern is shown in the likelihood surfaces depicted in Figure 5a-b. When the generating *ψ* is low (*ψ*=0.01 fossils/Myr; Fig. 5a), the likelihood surface includes the generating parameters. However, when *ψ* is increased by an order of magnitude (*ψ*=0.1 fossils/Myr; Fig. 5a-b), the likelihood surface begins to shift away from the generating parameters towards regions of parameter space represented by lower turnover rates and extinction fraction. While this downward bias in this particular example is somewhat subtle, this behavior is consistent across all models and all simulation scenarios examined here. It should be noted that it is possible that we have incorrectly interpreted the math described in Stadler et al. (2018). Admittedly, the implementation of this model was quite challenging, and is further complicated by the fact that FBD with stratigraphic ranges is not naturally nested within the canonical FBD model. That is, if all fossils are treated as single representatives of a stratigraphic range, the likelihood does not simply revert to the likelihood of the canonical FBD model. This is because of the assumption in the FBD with stratigraphic ranges that corrects for unobserved speciation occurring between two distinct stratigraphic ranges on the same edge (see Eq. 3). That being said, in the constant birth-death case, we implemented the model using the analytic equations presented in Stadler et al. (2018) as well as using the ordinary differential equations above and the likelihoods were identical across a range of parameter values.

**Figure 5.**
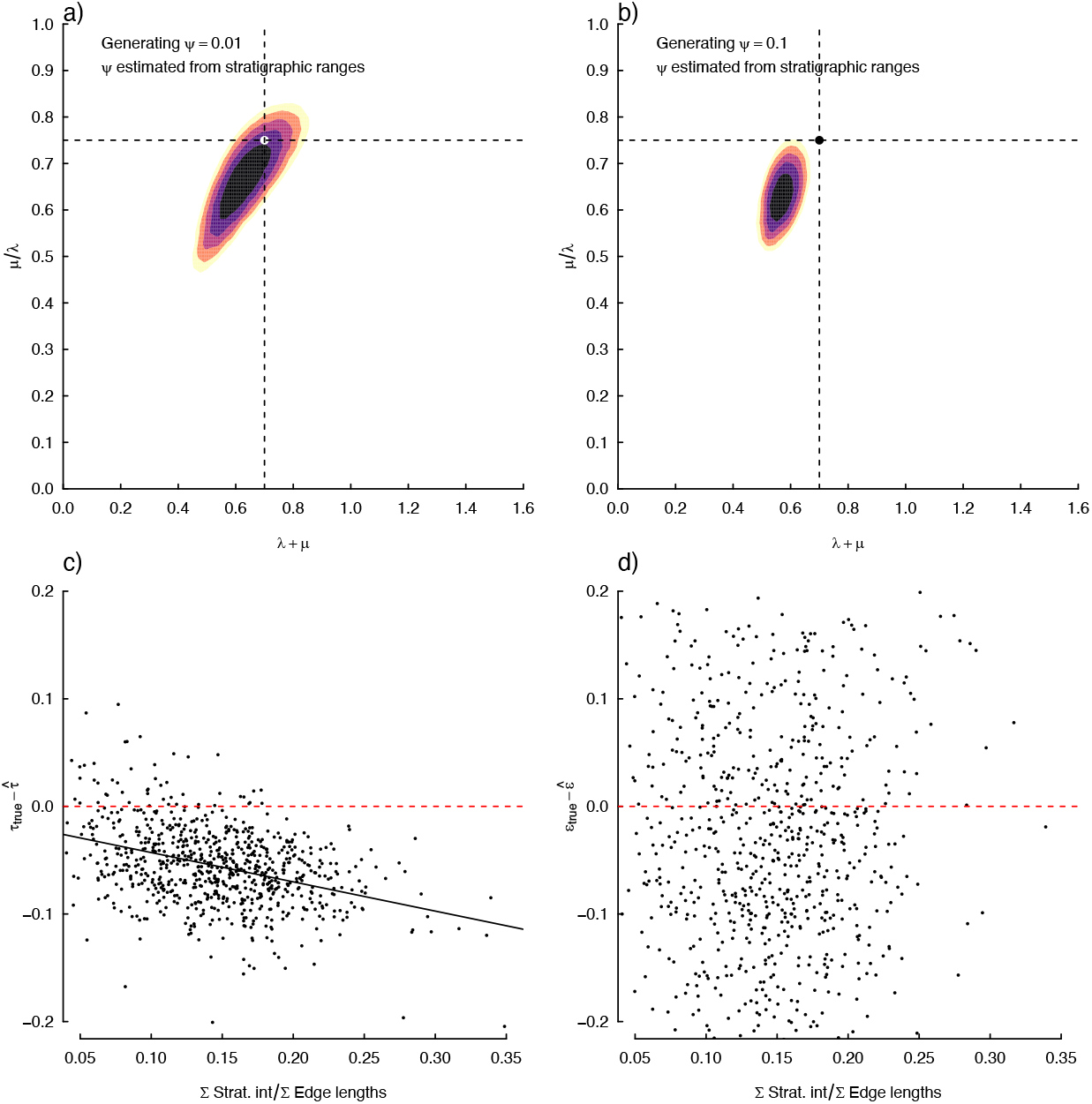
(a-b) Contour plots of the likelihood surface for the fossilized birth-death with stratigraphic ranges using the same tree and data set as shown in Figure 2. When the generating *ψ* is low, the likelihood surface includes the generating parameters, but when *ψ* is increased by an order of magnitude (*ψ*=0.1), the likelihood surface also begins to drift away from the generating parameters towards regions of parameter space represented by lower turnover rates and extinction fraction. (c) A stratigraphic range records the known duration of species on branches, and so we assume that no unobserved speciation or extinction takes place along these intervals. Since the same rates apply to all branches, and since turnover rate is a measure of the total number of speciation and extinction events, as the number of stratigraphic ranges and duration of each increase, turnover rates will increasingly be underestimated as the proportion of the stratigraphic ranges relative to the total tree length increases. Interestingly, this does not seem to impact estimates of extinction fraction (d).

In the absence of an implementation error, we suspect that the underestimation of turnover rate is likely due to the assumptions made when computing the probability of the portion of an edge representing a stratigraphic range. Under this model a stratigraphic range records the known duration of species on branches, and so we assume that no unobserved speciation or extinction takes place along these intervals. Since the same rates apply to all branches, and since turnover rate is a measure of the total number of speciation and extinction events, we might expect that as the number of stratigraphic ranges and duration of each increase, the rate estimates would likely reflect a balance between a large proportion of time in a tree where no unobserved speciation and extinction events have occurred. Thus, we would predict that the downward bias in turnover rates would be a function of the proportion of the tree consisting of stratigraphic ranges. We conducted a simulation where we evolved a set of 100 trees for 50 Myr, assuming the same generating rates as the ones that produced the trees in Figure 2. We also used a range of fossil sampling rates starting from *ψ*=0.05 fossils/Myr and increasing at 0.01 increments until reaching *ψ*=0.15 fossils/Myr. Consistent with our hypothesis, we found that as the proportion of the stratigraphic ranges relative to the total tree length increased (i.e., stratigraphic ranges accounted for increasingly more of the time represented in the tree), we found that this was significantly and negatively associated with an increased downward bias in turnover rates (*R*^2^=0.15, slope=-0.272, *P*<0.001). Interestingly, we did not find a similar trend with extinction fraction (*R*^2^<0.001, slope=0.09, *P*=0.246). While these results only speak to our implementation of the “symmetric speciation only” portion of Stadler et al. (2018; see Corollary 13), where only bifurcating speciation events are allowed, we suspect this behavior is general for all stratigraphic range models. Even if allowing for anagenetic and/or asymmetric speciation modes, we suspect turnover rate, which is a measure of the frequency of both speciation and extinction events, will be generally underestimated given all three speciation modes remove the possibility of an unobserved speciation and subsequent extinction event along a stratigraphic range edge.

## Concluding Remarks

Our results show that, if the trees fit the generating model well, then adding fossil taxa correctly might help with precision in rate estimations, but not with bias. However, for most cases if there is undersampling of fossils along branches (*k-*type fossils) the results are worse than ignoring fossils altogether, even when the placement of the sampled fossils was correct temporally and topologically. When placing the fossil set not as individual points on the tree, but rather as stratigraphic ranges, there is tendency for some measures of diversification to be systematically underestimated (i.e., turnover rate). All these results were obtained from a simulation framework where the diversification models used were far simpler than the simulated data. We also expected that at least the full fossil set would be able to somewhat rescue the results given that they were nearly three-times more numerous than extant species in certain scenarios (i.e., scenario 4). As we have emphasized elsewhere (see O’Meara and Beaulieu 2021), moving forward, focusing our comparisons of diversification rates at the tips of the tree remains a fruitful avenue of inquiry, even without fossils, given that most of the data from a phylogenetic tree of modern taxa exists nearer the present.

We hasten to emphasize that fossils remain key for understanding macroevolution — that is, important extinct groups like trilobites, sauropods, and extinct lycophytes are essential to our understanding of evolution in deep time. However, the idea that neontological studies of diversification are dramatically enhanced by sprinkling carefully chosen fossils on a tree and applying sophisticated fossilized birth-death models to understand rate variation does not seem supported by our simulations. At best, fossils have minor effects. At worst, they lead to less accurate inferences than removing them altogether. We even carefully biased our study to purposely give fossils the best possible chance, namely, by perfectly identifying each fossil, placing them with full certainty in a phylogeny, and assuming that their true sampling rate was constant through time. All this is impossible with empirical data, which is likely the product of variable fossilization rates (Wagner and Marcott 2013) that further complicates teasing apart variable extinction from variable preservation rates (see Foote et al. 2019; Louca et al. 2021). Taken together, our simulations provide us with little hope for the utility of fossils in real-world applications, at least with regards to estimating diversification rate heterogeneity across a tree.

One area where we suspect fossils will continue to be of great importance in neontological applications is in molecular divergence time analyses (e.g., Heath et al. 2014). Much of the information used in fitting most diversification models is the timing between events. So, if the relaxed clock does not properly account for clades with a faster rate of molecular substitution, most methods to smooth branches will yield ages that are much older than they should be (since the branches are still too long relative to the variation implicit in the clock; see Beaulieu et al. 2015), which will tend to also decrease the net diversification rate across the tree. However, having more fossils throughout the tree, we suspect, might alleviate this issue. It thus remains an open question (though with reasonable advocacy for various sides) whether methods that jointly estimate the topology, branching times, and diversification parameters using an FBD-type model give a better estimate of diversification rate parameters than alternate approaches (e.g., *r8s;* Sanderson 2002), which infer a chronogram agnostic to the diversification rate process and is subsequently used as input in diversification analysis programs. Our study simply demonstrates that in the case of a perfectly accurate (albeit perhaps undersampled) tree, fossils do not help substantially at estimating diversification process rates at the tips.

One motivation for this research, in the light of recent and well-earned concern about diversification methods using phylogenies of extant taxa (Kubo and Iwasa, 1995; Maddison and FitzJohn, 2015; Rabosky and Goldberg, 2015; Louca and Pennell, 2020; Louca et al. 2021) is to help guide empiricists seeking to create improvements in diversification rate estimation. There is understandable interest in fossils to improve diversification models. After all, we owe the discovery of extinction in the nineteenth century to fossils, and major events like mass extinctions or the Cambrian explosion were first and by far best known from purely the fossil record. While we did not investigate, and remain pessimistic, about the feasibility of estimating diversification processes through time, we were optimistic about the potential impact of fossils on rates at the tips as a first step. In our own reading of the foundational work of Stadler (2010), which was motivated by our own curiosity about a path for improvement, we were struck by the *m* versus *k* fossil distinction and what this might mean for its usability in practice, as well as how stratigraphic intervals might mitigate this issue. We were surprised that, at least for our particular use case, fossils provided at best little benefit to, and at worst harmed, inferences of diversification at the tips from trees of mostly modern taxa. This does not imply they may not help substantially in other areas of phylogenetics of extant taxa, but as a solution to this particular issue, they are less helpful than we hoped, and researchers might be more fruitful attempting other solutions first.

## Supplementary Materials

Data available from the following github repository: https://github.com/thej022214/Fossils_impact_BO

## Acknowledgements

We thank members of the Beaulieu and O’Meara labs for their comments and discussions of the ideas presented here.

